# Intuitive planning: global navigation through cognitive maps based on grid-like codes

**DOI:** 10.1101/421461

**Authors:** Alon B. Baram, Timothy H. Muller, James C.R. Whittington, Timothy E.J. Behrens

## Abstract

It is proposed that a cognitive map encoding the relationships between objects supports the ability to flexibly navigate the world. Place cells and grid cells provide evidence for such a map in a spatial context. Emerging evidence suggests analogous cells code for non-spatial information. Further, it has been shown that grid cells resemble the eigenvectors of the relationship between place cells and can be learnt from local inputs. Here we show that these locally-learnt eigenvectors contain not only local information but also global knowledge that can provide both distributions over future states as well as a global distance measure encoding approximate distances between every object in the world. By simply changing the weights in the grid cell population, it is possible to switch between computing these different measures. We demonstrate a simple algorithm can use these measures to globally navigate arbitrary topologies without searching more than one step ahead. We refer to this as intuitive planning.

## 1 Introduction

In order to generate complex behaviour that is flexible to changes in circumstance and internal goals, it has been argued that animals maintain a model of the relationships between objects in the world, often termed a cognitive map [25]. In physical space the neural substrate for such a map is hypothesized to exist in the hippocampal formation in the form of place cells [20] and grid cells [15]. Place cells are found in the hippocampus and fire when an animal is in a specific location, whereas grid cells, found in the entorhinal cortex, amongst other regions, fire in a periodic hexagonal lattice over the environment.

It is not yet clear why it is advantageous to encode such a map both locally (using place cells that fire in specific locations) and globally (using grid cells that each fire in many different locations, but whose population can encode location precisely). However, one attractive recent suggestion is that these grid codes can be used to efficiently solve navigation problems as much of the necessary computation is already solved in the representation. Angles and distances between different points in the map can be extracted from grid codes using inexpensive computations that do not require searches through place cells [5, 24], endowing the agent with a sense of direction for navigating the map intuitively.

Tolman’s original concept of a cognitive map was not, however, restricted to spatial relationships, but encompassed relationships that subserved flexible behaviour across multiple domains of life [25]. Indeed the hippocampal formation is critical for knowledge representation and generalisation in behavioural domains as distinct as social cognition [22], reward learning [14] and episodic memory [19]. Furthermore, neurons coding for non-spatial information in a fashion analogous to place cells have been found in human [21] and rodent [2] hippocampus, and signatures of grid-like activity have been reported in rodents encoding time rather than space [16] and in humans navigating conceptual, rather than spatial, knowledge [6].

It has recently been suggested that these different properties of hippocampal neurons can be reconciled formally if the idea of a place cell is generalised to encode a state in a reinforcement-learning world [23]. This conceptual insight allows the machinery of reinforcement learning theory to be applied to understand neural coding in the hippocampal formation. Indeed it has been shown that global navigational strategies can be performed in place cells, if the scope of their coding is broadened to predict future states using successor representations [8] that relate them to the navigational goal [23]. Furthermore, signatures of predictive state representations have been observed in human entorhinal cortex in discrete state space environments [13], and there is evidence that humans use predictive representations for planning [18].

Here, we present a different technique for navigating arbitrary conceptual worlds. This technique allows place cells to retain local scope but instead, as in the spatial domain [5, 24], relies on grid-like codes to generate globally optimal (or nearly optimal) routes with efficient computations that do not require any sequential search. Similar to space, this technique uses grid-like codes to endow the agent with an intuitive sense of direction, but in arbitrary relational worlds. We therefore refer to it as intuitive planning.

## 2 Global distances from local learning

We rely on the recent suggestions that grid cells can be understood as the eigenvectors of the relationship between place cells. This has been demonstrated both using a PCA of place cell firing [10], and using the eigenvectors of successor representations [23]. This view is consistent with the recently acknowledged importance of the feedback connections from place cells to grid cells [26, 4, 17].

The key insight in the current work is that grid cells representing eigenvectors learnt from only local inputs can be used to construct distributions of distant future states as well as a global distance matrix that encodes the approximate distances between every pair of points in the world. We are proposing this eigenstructure contains the information necessary to compute all possible distances without any search.

To see why this is true, consider the adjacency matrix, *A*, that defines relationships between neighbouring states (or place cells) by encoding the weight of the edge between *i* and *j*, as element *A*_*ij*_ (figure 1). To compute the distribution of states after *n* steps, it is necessary to compute the matrix *A*^*n*^. Notably, element 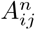 is the number of walks of length *n* from states *i* to *j*. As it is simply a power of *A*, all matrices *A*^*n*^ share the same eigenvectors of *A* (see figure 2 for example eigenvectors of *A* of a simple two-dimensional world):

**Figure 1:**
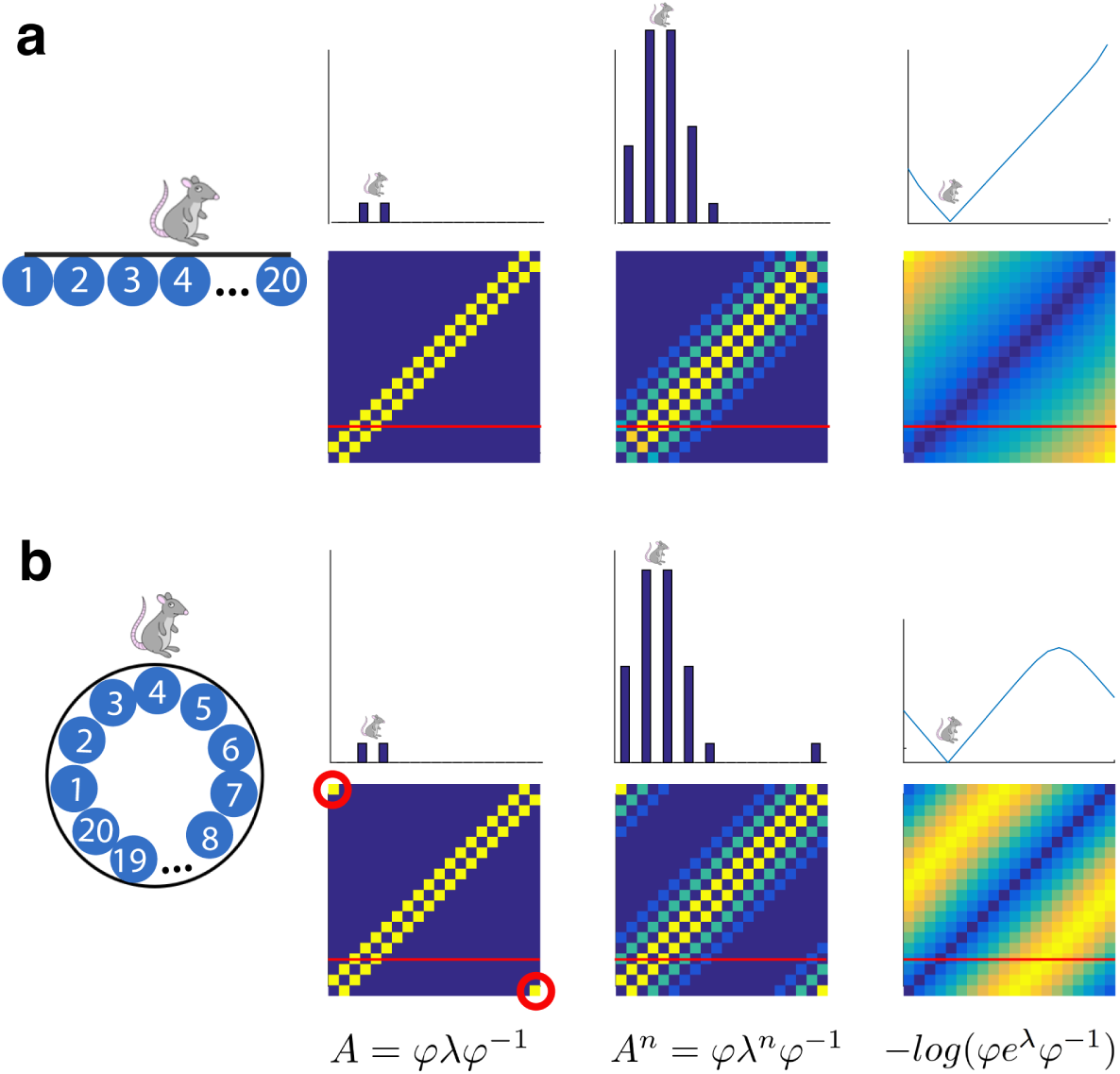
Construction of local and global knowledge from locally-learnt eigenvectors. **a)** A two-way linear track. For the three columns on the right, bottom is a matrix and top is a slice through that matrix (red line) at state four. From left to right: Schematic of the environment with a mouse occupying state four; the associated adjacency matrix *A*; *A*^*n*^, where *n* = 5, and therefore *A*_*ij*_ is the number of possible walks of length *n* from state *i* to state *j*;*−log*(*C*), where *C*_*ij*_ is the communicability from state *i* to state *j*. **b)** The same format as **a** for a two-way circular track. Red circles in the adjacency matrix mark the new adjacency between the ends of the linear track in **a** turning it in to a circular track. **Bottom**: Crucially, not only the matrix *A* containing local knowledge, but also *A*^*n*^ and *−log*(*C*) containing global knowledge can be constructed from the eigenvectors *ϕ* and eigenvalues *λ* of the matrix *A*. The local knowledge in state four is identical across the two environments (compare the slices through the adjacency matrices in **a** and **b**) but the global knowledge, obtained from the eigenvectors of *A*, is very different (compare slices in *A*^*n*^ and*−log*(*C*) across **a** and **b**).

**Figure 2:**
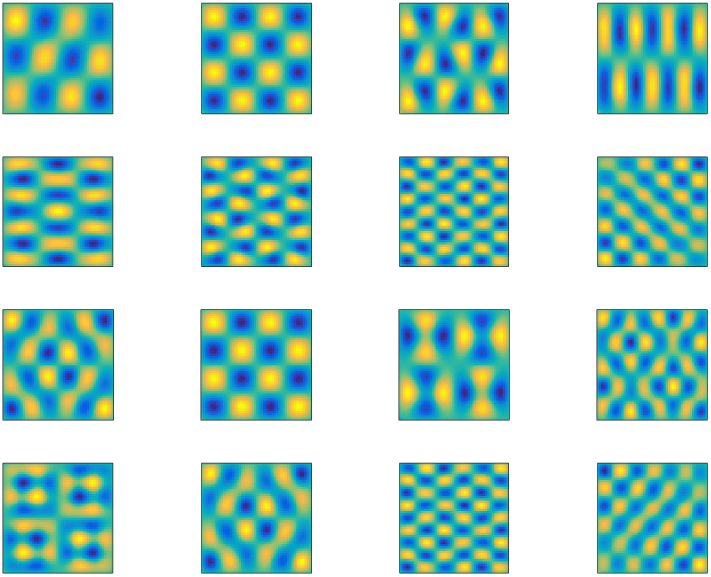
Example eigenvectors of the adjacency matrix of a simple two-dimensional world are periodic. Eigenvectors of the adjacency matrix *A* of a discrete-state two-dimensional box projected back in to the space of the box have periodic waveforms. These are similar to the eigenvectors reported in [10] before a non-negativity constraint was applied that rendered them hexagonal. The permitted transitions between states encoded in *A* were in the up/down and left/right directions.

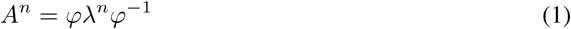

where the matrices *ϕ* and *λ* are the eigenvectors and eigenvalues of *A*, respectively.

Furthermore, any summation of *C*_*i*_ *A*^*ni*^, (where *n*_*i*_ is an integer, and *C*_*i*_ is an arbitrary scalar) will share these eigenvectors, so if we choose an approximate distance metric that is a weighted sum of future states, it is easily computable from the eigenvectors of *A*.

For example, consider the following two weighted sums:

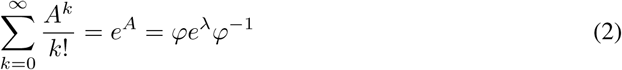

and

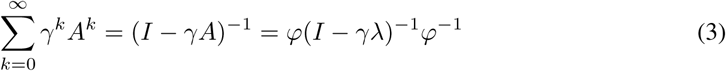

where 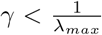 and *λ*_*max*_ is the largest eigenvalue of *A* in modulus. Throughout the paper, *γ* is set to the commonly used [3, 1] 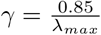. The infinite sums in equations 2 and 3 converge to the matrix exponential (*e*^*A*^) and resolvent ((*I − γA*)^*−*1^), respectively.

In each of these sums the dominant contribution to element (*i, j*) is given by the shortest path between *i* and *j*. Longer paths are dramatically downweighted (particularly with the matrix exponent, equation 2). Hence element (*i, j*) contains information that closely relates to the shortest distance between *i* and *j* (this is shown empirically below, including some exceptions). In graph theory, these two summations are used to compute the communicability between nodes of a graph [11, 1, 12]. The second of these two summations (equation 3) is similar to the successor representation in reinforcement learning [8], which is computed from the transition rather than adjacency matrix.

Notably, in order to compute these functions it is simply necessary to recombine the eigenvectors of *A* (grid cells) with different weights (figure 1). Instead of using the original eigenvalues *λ, λ*^*n*^ will give the distribution of future states after *n* steps, *e*^*λ*^ will give the matrix exponent, and (*I−γλ*)^−1^ will give the resolvent (or successor representation). Further, these reweightings are inexpensive to implement because the number of eigenvalues is equal to the number of grid cells (eigenvectors), not the number of distances (relationships) that can be reconstructed.

We emphasise that these measures work for arbitrary topology environments. They accurately reflect communicability in environments that cannot be easily mapped on to normal maps, such as those containing wormholes linking disparate parts of a map (figure 3), as well as environments with one-way links between states, which result in complex eigenvectors and eigenvalues (figure 3). Such arbitrary topologies are more likely those found in non-spatial cognitive maps.

**Figure 3:**
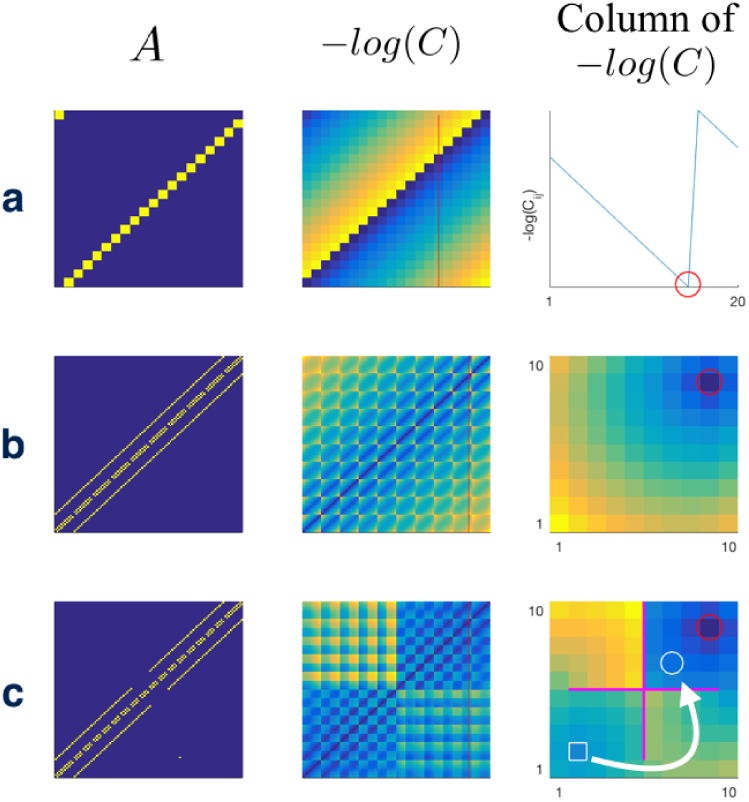
Communicability measures in arbitrary topologies. **a)** A one-way linear track. Left: The adjacency matrix *A*. Middle: the matrix *−log*(*C*) where element *C*_*ij*_ is the communicability between states *i* and *j*. Red line marks the column of a target state (note this is different to figure 1 where it is the row of the current state). Right: This column represented on the space defined by *A*. Red circle indicates target state. **b)** A two dimensional 10×10 box. The same format as **a** except the target column of *−log*(*C*) is represented as a heatmap on a two-dimensional environment with warmer colours corresponding to higher values. **c)** A box with barriers and a wormhole connecting states which are not adjacent in a two-dimensional space. Same format as **b**. States separated by a magenta line are not adjacent. White squares and circles mark entrance and exit gates of a wormhole, respectively: the states are adjacent yet distant in the regular two-dimensional topology. The conventions used in this figure will be used throughout the paper.

Both the matrix exponential and resolvent measures can be approximately linearly related to the true distance, as defined by the length of the shortest path between states, by taking their *−log* (figure 4), a measure we refer to as intuitive distance. This relationship is dependent on the sets of walks between pairs of states with the same distance. In some environments, the set of walks between a pair of states can be obtained by a permutation from the set of walks between another pair of states with the same true distance. In such a case there is no variance in the relationship between intuitive and true distance (figure 4a). However, in environments where there are many short, yet not necessarily shortest, paths between states, the communicability might differ between pairs of states with the same true distance, and hence so will the intuitive distance (figure 4b). This predicts failures to navigate the shortest path between states (see below).

**Figure 4:**
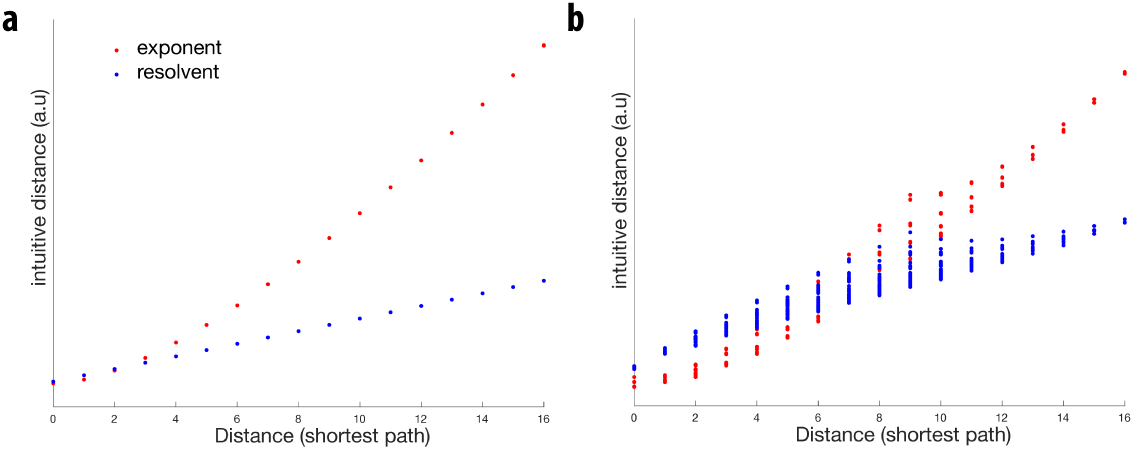
Approximately linear relationship between intuitive distance and true distance. Intuitive distances between pairs of states are approximately linearly related to the true distance, as defined by the length of the shortest path. **a)** Intuitive distance reliably increases with true distance on a two way, one-dimensional track (the same as that reported in figure 1b). **b)** Spread of intuitive distances for a given true distance in a 10×10 two-dimensional box (the same as that reported in figure 3b), because communicability is influenced by the number of routes between states.

## 3 Global navigation from local computation

Given current and target states, as well as the eigenvectors and eigenvalues, all calculations necessary for navigation are local: The agent needs to calculate which of the available next states has the shortest distance to the target. This can be done in the case of the matrix exponential by calculating for target state *T* and each neighbouring state, *N* :

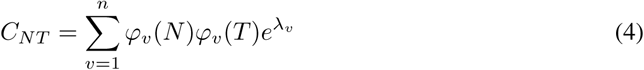

and in the case of the resolvent by calculating:

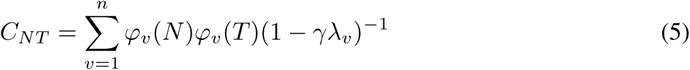

where *ϕ*_*v*_ (*N*) is the *N* th element of the *v*th eigenvector of the adjacency matrix associated with the eigenvalue *λ*_*v*_ and *n* is the number of eigenvectors. The agent can then proceed to the neighbouring state with the maximal *C*_*NT*_, or equivalently minimal intuitive distance *−log*(*C*_*NT*_). This procedure finds trajectories that are approximately globally optimal without any global (tree) search. Because it stores the relational structure, the algorithm can flexibly reroute when the values of candidate target states change, for example due to changing internal goals. Further, parallel neural circuitry can naturally implement the summations in equations 4 and 5.

## 4 Navigation using intuitive planning

We constructed several discrete-state mazes and tested whether intuitive planning could successfully navigate between start and target locations. We demonstrate the algorithm successfully navigates the shortest route to the target in several two-dimensional, undirected mazes (figure 5). Cognitive maps encoding non-spatial relationships, however, will likely have arbitrary topologies. We show that the algorithm generalises to arbitrary topologies as its performance is not affected by the insertion of a wormhole in to the maze, rendering it no longer topologically two-dimensional (figure 5). Successful navigation of the shortest route to the target, as defined by the minimum number of transitions to reach it, is achieved in all of the above mazes when using both the matrix exponential and resolvent measures. The mazes in figure 5 can be mapped on to two-dimensional spaces for visualisation purposes. The algorithm also achieved successful navigation to the target in randomly generated, unweighted graphs, provided there was a route to the target (figure 5).

**Figure 5:**
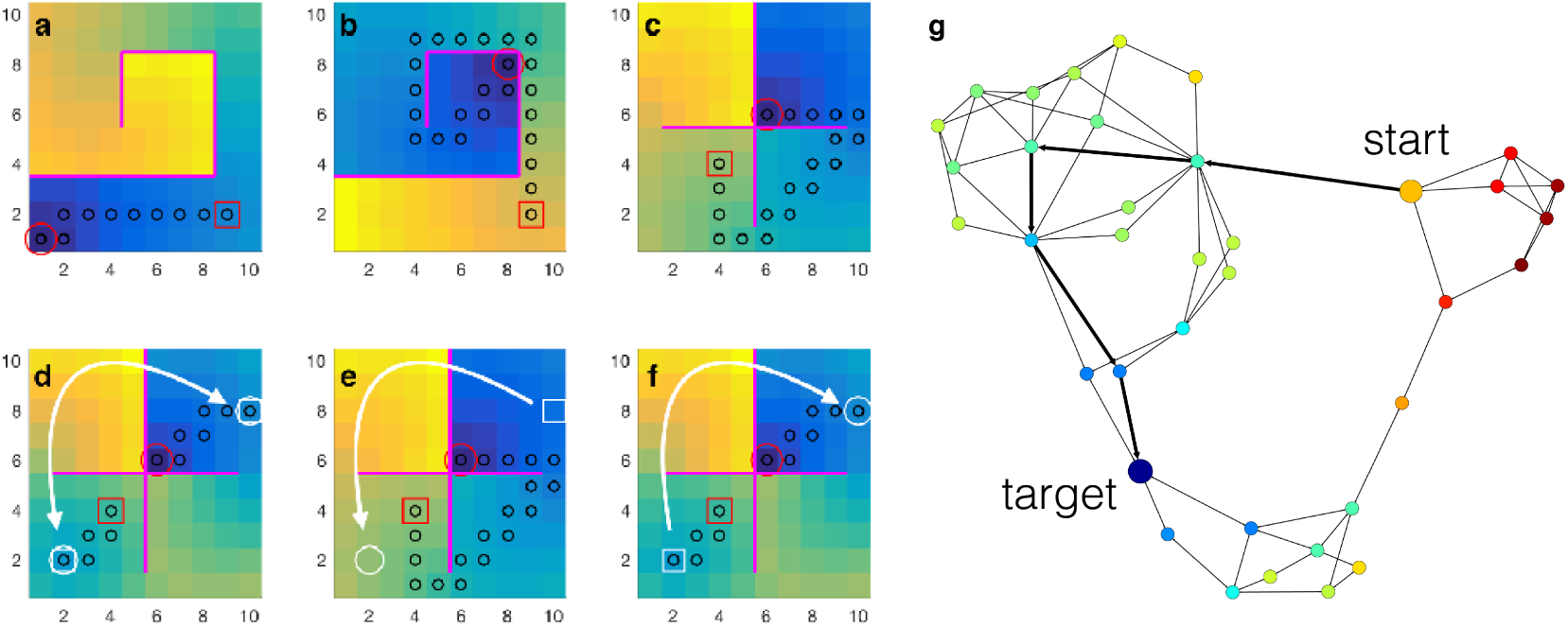
Navigation in arbitrary topology worlds. Successful navigation from start (red square) to target (red circle) states in two-dimensional worlds with barriers and differing start and target states (**a-c**). Introducing wormholes that connect disparate parts of the map and therefore warp it rendering it no longer described by a simple map does not affect the ability to successfully navigate to the target (**d-f**). Nor does the introduction of a one-way wormhole (**e-f**), resulting in complex eigenvalues and eigenvectors. In all navigation simulations, state transitions were permitted in the up/down and left/right directions (unless, for example, blocked by a barrier), and black circles denote the agent’s path. **g**) Successful navigation in a randomly generated graph.

Since the relationship between intuitive distance and true distance is only approximately linear (figure 4), it is possible to construct mazes where the algorithm successfully navigates to the target, but does not take the shortest route. This is achieved by increasing the communicability of initial states not associated with the shortest route, but with multiple longer routes to the target (figure 6).

**Figure 6:**
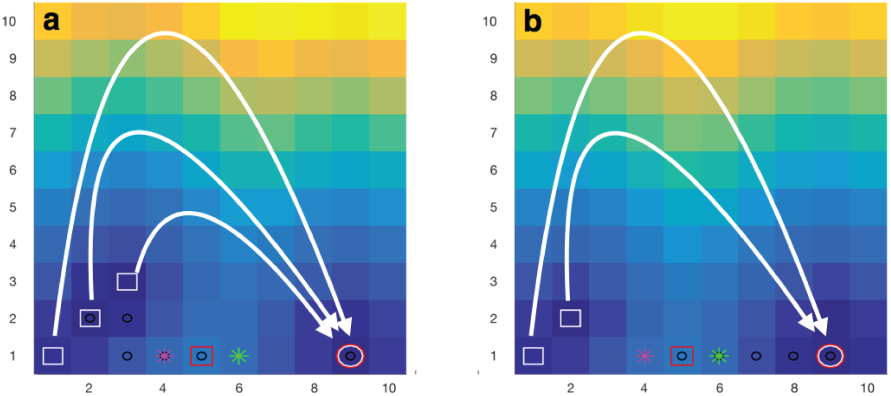
Failure to navigate the shortest path. Introducing multiple wormholes that cause multiple longer routes to the target to pass through a nearby state (magenta star) that is not in the direction of the shortest route (green star) causes the agent to take a longer route to the target (**a**). With fewer such wormholes, the shortest route is taken (**b**)

## 5 Experimental predictions

This theoretical work makes some experimental predictions:

- Unlike explicit planning [9], such intuitive planning will not require a precession through place cells leading to a goal.
- In intuitive planning the distance to a goal does not influence the difficulty of the next decision. The opposite is true in explicit planning (tree-search). We therefore predict that for local goals, distance to goal (where explicit planning is possible) will predict reaction time, but for distant goals it will not.
- The intuitive distance measures are most likely to differ from shortest-path measure when there are many possible routes to the goal that pass through one nearby state, but where the shortest route passes through a different state (figure 6). This predicts a precise pattern of planning errors.

## 6 Discussion

We believe that the observation that grid cell patterns can be related to the eigenspectrum of local place cell transitions [10, 23] is an important one. We have shown that one consequence of this observation is that grid cell patterns contain information not only about the current location but also about all possible future locations. It is therefore possible to compute distance metrics between every pair of locations in the state space using grid patterns that have been learnt from only one-step transitions. With one choice of weighting of future states, this distance metric allows rapid computation of successor-like representations [8, 23] but with states (place cells) that retain local scope. Other distance metrics can also be computed. Like successor representations, this provides only a partial solution to the model-based reinforcement-learning problem leaving open the problem of how values are assigned to different target locations. The eigenvector-based computations suggested here are easily implemented in parallel, and therefore suggest natural implementations in recurrent neural circuits that support grid codes. Indeed, Corneil & Gerstner [7] have elegantly implemented a planning algorithm based on the eigendecomposition of the SR of the environment in a recurrent attractor neural network. The global metrics we have presented here allow global navigation through arbitrary topology maps without searching more than one step ahead. We refer to this as intuitive planning. Such a strategy could be combined, for example, with hierarchical representations of state space to further reduce computational complexity. Together, such strategies demonstrate how the expensive computations that underlie state-space navigation can be finessed by efficient choices of neuronal representation.

